# Scaling genetic discovery through automated post-GWAS interpretation

**DOI:** 10.64898/2026.10.23.761278

**Authors:** Libo Qiao, Shuang Gu, Ye Yang, Yingnan Han, Xuanzhao Li, Xing Wan, Xiwen Zhang, Qinhao Zhang, Tingting Hu, Yan Zhang, Faming Chen, Jiaheng He, Jingxuan Wang, Zhenyu Tang, Peng Ye, Lilong Wang, Erping Long, Xiaosong Wang, Lei Bai, Fulong Yu

**Affiliations:** GMU-GIBH Joint School of Life Sciences, The Guangdong-Hong Kong-Macao Joint Laboratory for Cell Fate Regulation and Diseases, Guangzhou Medical University, Guangzhou, Guangdong, China; Guangzhou National Laboratory, Guangzhou, Guangdong, China; State Key Laboratory of Respiratory Disease, Guangzhou, Guangdong, China; Shanghai Artificial Intelligence Laboratory, Shanghai, China; Medical Research Institute, Guangdong Provincial People’s Hospital (Guangdong Academy of Medical Sciences), Southern Medical University, Guangzhou, Guangdong, China; Institute of Basic Medical Sciences, Chinese Academy of Medical Sciences and Peking Union Medical College, Beijing, China

## Abstract

Genome-wide association studies (GWAS) have made association discovery systematic and cumulative, yet evidence for interpreting associated loci remains selective, heterogeneous and difficult to accumulate across studies. Here we present Post-GWAS Intelligence (PGI), which couples context-dependent post-GWAS analysis to a common representation of analysis-derived evidence. Its multi-agent engine, VariantAgent, executes workflows, records tool-derived results in standardized variant-centred units and generates trait-level interpretations linked to their analytical basis and evidential status. Complementary benchmarks showed that VariantAgent outperformed tested general-purpose and biomedical agents, with its largest advantage when success required both a correct answer and a valid analytical trace. Across 34 published GWAS, VariantAgent achieved a mean per-study recovery of 82.1% for directly comparable findings. The largest gains lay downstream of association mapping, where uniform analysis extended published interpretations and identified regulatory relationships at established risk loci not previously reported in the source studies. Analysis of 1,041 GWAS yielded 530,949 evidence units and corresponding reports, which PGI organized into a cross-trait catalogue. Catalogue interrogation resolved an apparent cross-trait overlap into distinct signals and trait-specific biological hypotheses. This work demonstrates how PGI can extend genetic discovery beyond association mapping by enabling biological evidence to accumulate across studies.

## INTRODUCTION

Twenty-five years after publication of the draft human genome sequence, genetic association discovery has become systematic and cumulative^1^. The reference genome established a common coordinate framework for locating and comparing variation across studies. Advances in genotyping, sequencing and biobank-scale phenotyping then enabled genome-wide association studies (GWAS) to map genotype-phenotype associations across millions of individuals and a growing range of traits^2–4^. The NHGRI-EBI GWAS Catalog now contains more than one million curated variant-trait associations reported in the literature^5^. Yet an associated locus marks the beginning rather than the completion of genetic discovery. It does not by itself identify the causal variants, effector genes, relevant cellular contexts or mechanisms through which genetic variation influences phenotype. Resolving these links is what allows human genetics to yield biological understanding and inform intervention^6^. Association signals accumulate in shared catalogues, but the evidence used to interpret them remains fragmented across studies.

Post-GWAS analysis extends beyond annotation. Fine-mapping distinguishes among correlated variants. Molecular QTLs and chromatin maps connect variants to genes and cellular contexts, sequence and structure models predict molecular consequences, and perturbation experiments test selected links^6,7^. These approaches differ in what they measure and in how strongly they support a mechanism. Their relevance also depends on ancestry, linkage disequilibrium, variant class, available data and biological context. A valid interpretation must therefore retain whether each link was observed, statistically supported, predicted or inferred. The task is to connect these layers into a defensible account of how an association may act without erasing those distinctions. Studies combining fine-mapping, cell-resolved regulatory maps and targeted perturbation can resolve individual loci in considerable mechanistic depth^8–10^. Such analyses, however, are usually designed around one trait or locus and require extensive specialist judgement. The challenge is to extend this rigour across studies without treating predictions as equivalent to measurements.

The growing abundance of analytical methods and functional data has exposed a broader bottleneck. Predefined pipelines can run selected post-GWAS analyses consistently, whereas resources such as the IGVF Catalog make variant-effect measurements and predictions accessible for reuse^11,12^. Making analyses repeatable and evidence retrievable does not make interpretation cumulative. Pipelines are designed around predefined workflows, and functional resources do not by themselves determine which evidence is applicable to a particular GWAS or how it should be combined with results generated during analysis^13,14^. Biomedical agents can connect these components more flexibly^15^, but flexibility does not guarantee analytical validity. Genome build, allele alignment, ancestry-matched linkage disequilibrium, statistical assumptions and biological context can each alter the downstream conclusion^16^. A fluent report can still conceal invalid analytical choices and conflate analytical results, retrieved evidence and model-generated inference. Methodological choices and intermediate outputs often remain dispersed across scripts and supplementary files, while final interpretations are compressed into prose. Reliable automation therefore requires the analytical record to be structured as it is generated. Tool-derived results must remain distinct from the interpretation built from them, so that claims can be checked, compared across studies and revised as evidence changes^17^.

Here we establish PGI, in which VariantAgent performs domain-guided analyses and records tool-derived results in standardized variant-centred units that remain separate from, but linked to, trait-level reports. In matched reanalyses of 34 published GWAS, VariantAgent recovered a mean of 82.1% of directly comparable findings. Uniform reanalysis revealed a marked downstream asymmetry in coverage. Independent signals and prioritized variants increased several-fold, whereas prioritized genes, mechanistic hypotheses and perturbation evidence increased by more than an order of magnitude. Across 1,041 GWAS, VariantAgent generated 530,949 evidence units together with trait-level reports, which PGI organized for comparison across studies and traits. These results indicate that downstream evidence remains unevenly interrogated and difficult to accumulate. PGI thereby extends the cumulative logic of GWAS from association signals to the evidence used to interpret them.

## RESULTS

### PGI enables systematic and traceable post-GWAS interpretation

We developed Post-GWAS Intelligence (PGI) to support context-dependent post-GWAS analysis while standardizing the evidence generated across studies (Fig. 1a). PGI connects VariantAgent, a supervised multi-agent analytical engine, to a shared evidence architecture. Each run produces standardized variant-centred evidence units and evidence-grounded trait-level reports, retaining study context, provenance and evidential status. The evidence units distinguish statistical associations, source-backed molecular relationships, computational predictions and inferred candidates. PGI indexes these outputs across studies to build an analysis-derived catalogue, distinct from the biomedical knowledge graph used during execution.

**Figure 1.**
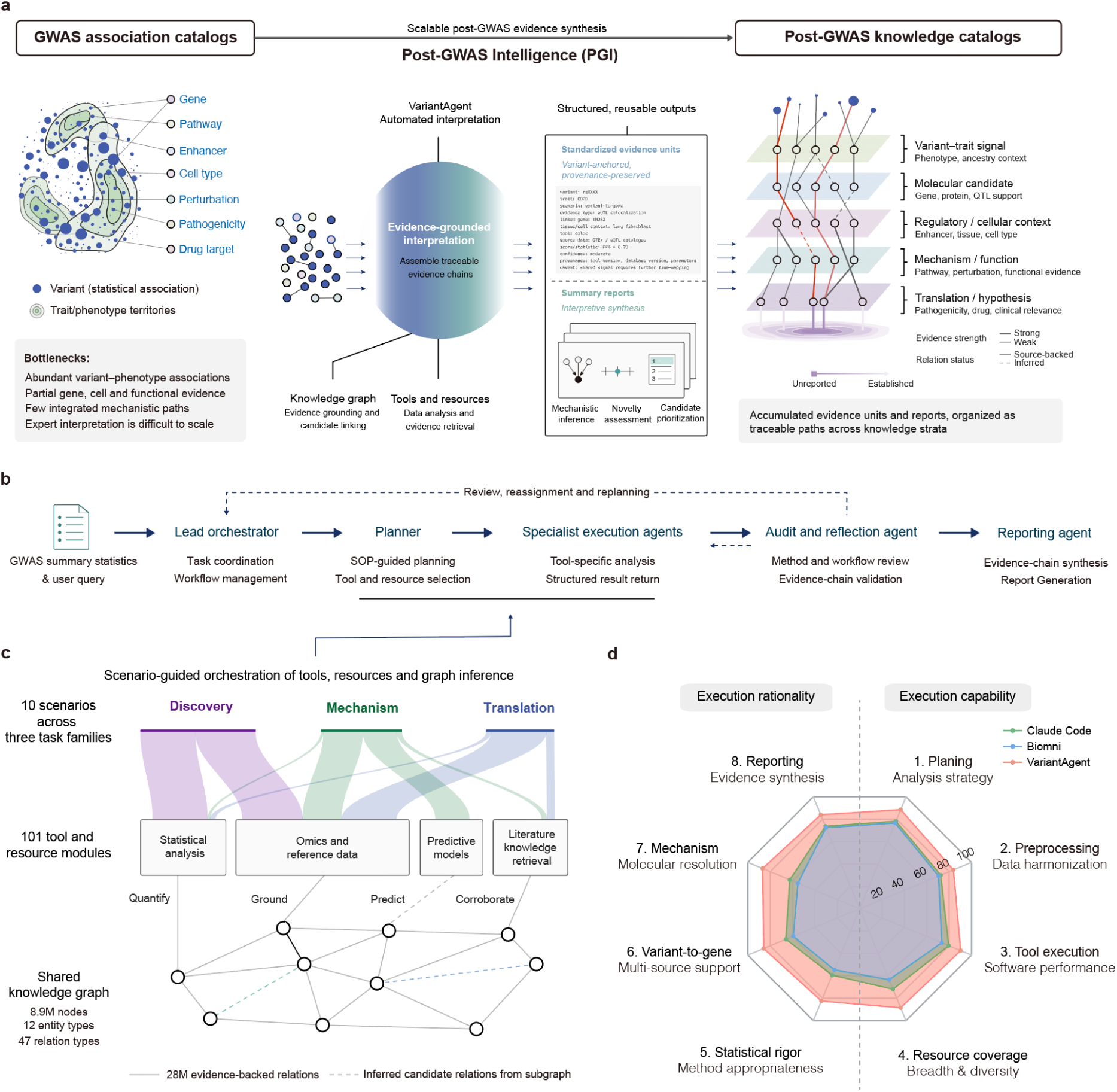
Architecture, analytical scope and process-level evaluation of Post-GWAS Intelligence. **a,** Conceptual organization of PGI. GWAS association catalogues provide variant–trait signals, whereas PGI connects domain-guided analysis to structured evidence accumulation. Its execution engine, VariantAgent, combines analytical tools and resources with a biomedical knowledge graph to assemble supported paths from association signals to molecular candidates, regulatory and cellular contexts, biological mechanisms and translational hypotheses. Each run retains an execution record and produces two linked scientific outputs. Standardized variant-centred evidence units preserve tool-derived and retrieved records together with their study context, provenance and evidential status. Evidence-grounded trait-level reports use these units for candidate prioritization, mechanistic interpretation and hypothesis generation. PGI indexes the units and reports across studies and traits to form an analysis-derived post-GWAS knowledge catalogue. Source-backed and inferred relationships remain separately labelled. The biomedical knowledge graph used during execution is distinct from the resulting PGI catalogue. **b,** Supervised multi-agent execution in VariantAgent. A lead orchestrator acts as the team leader, retaining the analytical objective and workflow state throughout an analysis. A planner converts assigned tasks into SOP-guided plans, and a dynamically assembled team of specialist execution agents operates the tools and resources summarized in **c**. The number and composition of these agents vary with the scientific question and available data. A reporting agent synthesizes accepted evidence, whereas an audit and reflection agent returns workflow-level feedback to the lead orchestrator for targeted reassignment, replanning or re-execution. **c,** Scenario-guided orchestration of analytical capabilities. Ten post-GWAS scenarios spanning discovery, mechanism and translation are mapped to 101 tool and resource modules covering statistical analysis, omics and reference data, predictive models, and literature or knowledge retrieval. A shared biomedical knowledge graph containing approximately 8.9 million nodes, 12 entity types, 47 relation types and 28 million source-backed relationships supports entity alignment, evidence retrieval and candidate linking. Candidate relationships nominated from graph paths remain explicitly distinguished from source-backed relationships. **d,** Process-level comparison of VariantAgent, Claude Code and Biomni across eight prespecified dimensions in ten full-process GWAS evaluations. Analytical execution comprised resource awareness and planning, preprocessing and cross-resource harmonization, software and tool execution, and resource coverage and diversity. Methodological validity comprised statistical appropriateness, multi-source variant-to-gene support, mechanistic evidence integration, and evidence-grounded reporting and hypothesis generation. Points show normalized rubric-based scores on a scale from 0 to 100, with higher values indicating stronger performance. Evaluation and score aggregation are described in Methods.

VariantAgent coordinates SOP-guided planning and specialist execution through a supervised multi-agent architecture (Fig. 1b). A lead orchestrator maintains the analytical objective and workflow state, while audit and reflection support methodological checks and targeted re-execution. Ten analytical scenarios span post-GWAS discovery, mechanism interpretation and translational evidence (Fig. 1c). The GWAS-KG biomedical knowledge graph supports evidence retrieval and candidate linking; source-backed and graph-inferred relationships remain distinguished.

To evaluate analytical reliability, we assessed VariantAgent on ten GWAS workflows across eight prespecified dimensions covering execution and methodological validity (Fig. 1d). VariantAgent achieved an overall full-process score of 86.3, compared with 70.4 for Claude Code and 65.7 for Biomni, and ranked first across all ten workflows. Three geneticists blinded to system identity evaluated the outputs; expert and agent scores showed high agreement (ICC = 0.91). VariantAgent also showed greater consistency across repeated runs (Extended Data Fig. 1a). Under matched foundation-model, tool, resource and execution-budget conditions, removing scenario guidance and workflow-level audit reduced the overall score from 88.4 to 79.8 (Extended Data Fig. 1b), supporting the contribution of domain-guided execution and methodological review.

### VariantAgent improves analytical accuracy and reliability

We evaluated VariantAgent using two complementary answer-level benchmarks (Fig. 2). The first comprised 313 published genetic-reasoning questions from three benchmarks^15,18,19^ spanning phenotype–gene, variant–phenotype and variant–gene relationships; the second comprised 26 questions requiring analysis of supplied GWAS summary statistics from ten datasets^20,22–31^. The GWAS-grounded benchmark’s construction and execution are detailed in Extended Data Fig. 2; these evaluations were distinct from the ten-workflow full-process assessment in Fig. 1.

**Figure 2.**
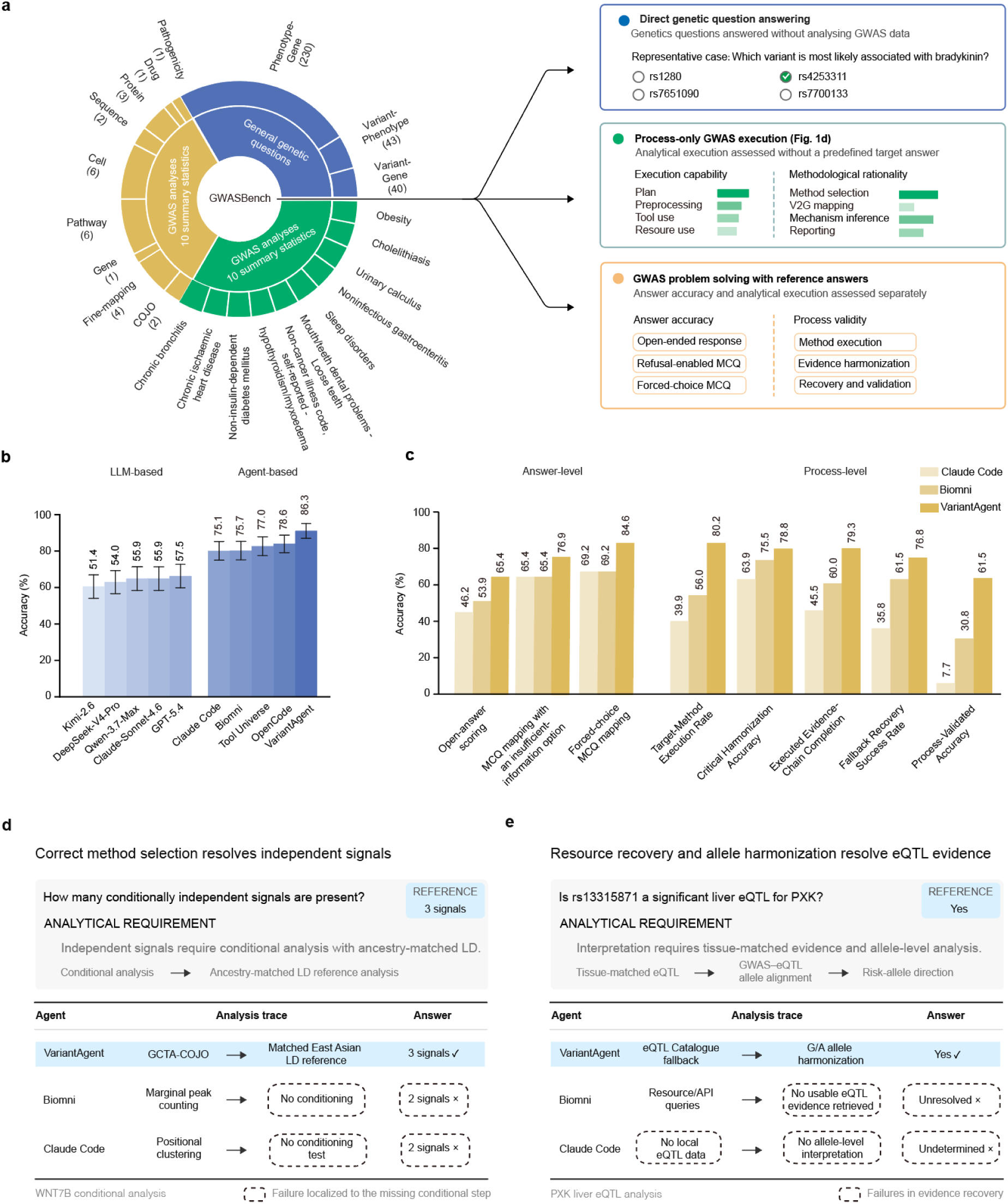
Benchmarking reveals a gap between answer accuracy and process-valid post-GWAS analysis. **a,** Overview of three complementary evaluation settings. Three published genetic-reasoning benchmarks contributed 313 questions, including 230 phenotype-to-gene, 43 variant-to-phenotype and 40 variant-to-gene questions. A separate set of ten GWAS datasets was used for the eight-dimensional full-workflow evaluation reported in Fig. 1d. Ten additional GWAS summary-statistic datasets were used to construct 26 study-specific questions covering conditional association analysis, fine-mapping, gene and pathway prioritization, cellular context, molecular consequence and translational evidence. The last setting was evaluated at both answer and execution-trace levels. **b,** Accuracy across the 313 published genetic-reasoning questions. Values indicate the number and percentage of questions answered correctly; error bars show 95% confidence intervals calculated by nonparametric bootstrap resampling at the question level (10,000 bootstrap replicates; percentile method). Relationship-class results are shown in Extended Data Fig. 3. **c,** Answer- and process-level performance on the 26 GWAS-grounded questions. Answer accuracy was evaluated under open-answer scoring and two alternative multiple-choice mappings, with or without an explicit insufficient-information option. Process-level performance was evaluated by target-method execution, critical harmonization, executed evidence-chain completion, fallback recovery and process-validated accuracy. **d,** Representative conditional-analysis case at *WNT7B*. VariantAgent recovered the three reference signals using GCTA-COJO with ancestry-matched LD, whereas Biomni and Claude Code omitted the conditioning step and returned two signals. **e,** Representative liver eQTL case for *PXK*. VariantAgent recovered alternative eQTL evidence and completed allele harmonization, whereas Biomni and Claude Code failed to resolve the question because evidence retrieval or allele-level interpretation was incomplete.

Across the 313 published questions, VariantAgent answered 270 correctly (86.3%), compared with 246 (78.6%) for OpenCode, 241 (77.0%) for Tool Universe^21^ and 235 (75.1%) for Claude Code (Fig. 2b, Extended Data Fig. 3). Its advantage over the strongest comparator was 7.7 percentage points (95% CI, 3.8–11.8), based on 10,000 paired bootstrap resamples. On the 26 GWAS-grounded questions, VariantAgent achieved 65.4% (17/26) open-answer accuracy, compared with 53.9% (14/26) for Biomni and 46.2% (12/26) for Claude Code (Fig. 2c). These percentages describe separate benchmarks and should not be interpreted as directly comparable measures.

We further audited the 26 GWAS-grounded execution traces to distinguish correct answers from methodologically supported answers. Process-validated accuracy was 61.5% for VariantAgent, 30.8% for Biomni and 7.7% for Claude Code. Among correct answers, 94.1%, 57.1% and 16.7%, respectively, had valid supporting traces. VariantAgent also more consistently executed required tools, completed evidence chains and recovered from unavailable resources (Fig. 2d,e). Trace inspection illustrated the importance of ancestry-matched conditional analysis at the WNT7B locus and explicit fallback handling for PXK eQTL evidence. VariantAgent’s evidence-admission decisions agreed with blinded expert adjudication in 92.3% of traces. Together, these results show that answer correctness alone can obscure differences in analytical validity.

### VariantAgent recapitulates T1D genetic architecture and extends mechanistic evidence

To assess whether automated post-GWAS analysis could reproduce expert-led biological interpretation, we applied VariantAgent to a European-ancestry type 1 diabetes (T1D) GWAS^22^ comprising 18,942 cases and 501,638 controls (Fig. 3). GCTA-COJO identified 324 independent signals, including 125 that reached genome-wide significance in both marginal and conditional analyses (Fig. 3a). These signals encompassed established T1D loci, including PTPN22, IFIH1, CTLA4, IL2RA and TYK2. Fine-mapping with ABF, SuSiE and FINEMAP prioritized 910 candidate variants supported by at least two methods, including 52 supported by all three (Fig. 3b).

**Figure 3.**
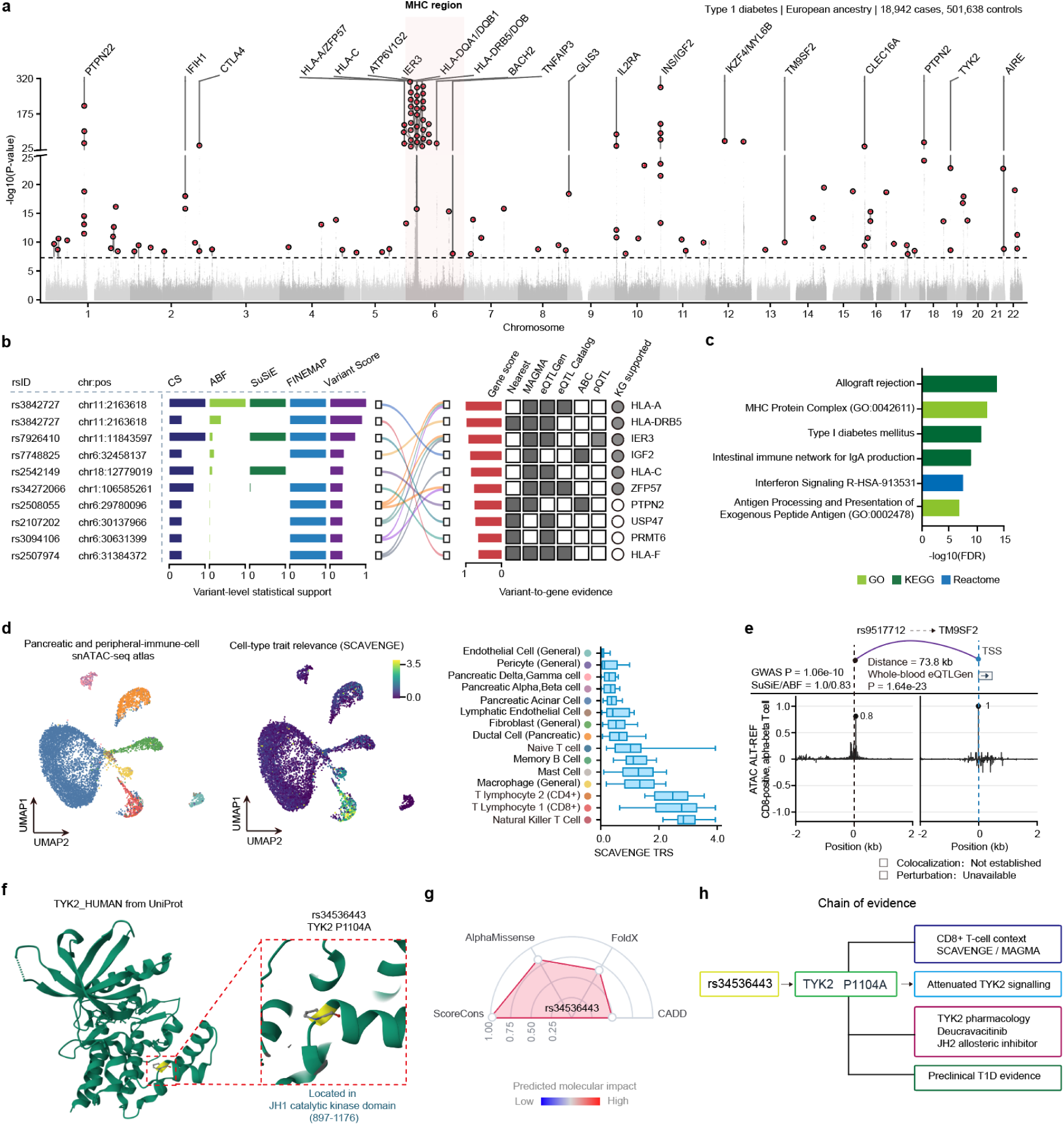
Full-process analysis of type 1 diabetes distinguishes candidates from functionally anchored evidence paths. **a,** Marginal association landscape for type 1 diabetes in a European-ancestry dataset comprising 18,942 cases and 501,638 controls. GCTA–COJO was performed using 1000 Genomes Project European GRCh38 PLINK reference panel. Red circles denote 125 signals reaching P < 5 × 10−8 in both the marginal and conditional analyses. Orange outlined circles denote 199 additional COJO-selected signals that did not reach genome-wide significance in the marginal scan. The extended MHC region is shaded. The y axis shows marginal GWAS −log_10_(*P*). **b,** Cross-model variant prioritization and variant-to-gene evidence. At an operational method-specific posterior threshold of 0.1, 910 unique variants were prioritized by at least two of ABF, SuSiE and FINEMAP, including 52 prioritized by all three. Model-specific posterior probabilities and credible-set membership are displayed separately. Representative variants and genes were selected according to gene-first ranking by cross-source normalized PIP-weighted variant-to-gene support. Variant-to-gene evidence includes proximity, molecular QTLs, activity-by-contact links and pQTLs. MAGMA gene-level association is shown separately, and graph-nominated candidate links are distinguished from source-backed relationships. Representative variants were then selected for each displayed gene by prioritizing source-backed variant-gene links with stronger fine-mapping posterior support and higher variant score. **c,** Enrichment of prioritized genes across non-redundant immune and regulatory pathways. Enrichment was tested using Enrichr over-representation analysis against the default background of each Enrichr gene-set library and adjusted using Benjamini-Hochberg FDR correction as reported by Enrichr. Results from the primary and MHC-excluded analyses are shown. **d,** Cellular distribution of T1D trait-relevance scores in the pancreatic and peripheral-immune single-nucleus ATAC-seq datasets. UMAPs show cell annotations and SCAVENGE trait-relevance scores. These scores provide trait-level context and are not assigned to individual loci. UMAP, uniform manifold approximation and projection. **e,** Candidate evidence path linking rs9517712 to *TM9SF2*. Fine-mapping support, genomic distance, whole-blood eQTL association and the AlphaGenome-predicted ref–alt difference in ATAC-seq track in CD8-positive alpha-beta T cells (CL:0000625; AlphaGenome ALT-minus-REF delta) are shown. The variant-to-gene relationship is displayed as a candidate link because colocalization, a physical enhancer–promoter connection and perturbational validation have not been established. **f,** Structural localization of the TYK2 P1104A substitution encoded by rs34536443 using UniProt P29597. P1104A lies within the JH1 catalytic domain. **g,** Molecular annotations of P1104A. Raw AlphaMissense, FoldX, CADD and conservation values are displayed separately with their published interpretation thresholds. The scores are not combined because they quantify different molecular properties. **h,** Branched evidence representation for rs34536443 and *TYK2*. The central path links the protective T1D association to P1104A and published evidence for context-dependent attenuation of *TYK2* signalling. Trait-level T-cell relevance, pharmacological targeting of the JH2 domain by deucravacitinib and T1D-relevant preclinical studies are shown as separate contextual branches. Solid edges denote directly observed or source-backed relationships, dashed edges denote predicted or inferred relationships, and grey edges denote trait-level context.

Integration of genetic, molecular QTL and regulatory evidence recovered established T1D genes and highlighted immune-related mechanisms. Gene-set analysis identified antigen presentation, interferon signalling and autoimmune pathways (Fig. 3c), while SCAVENGE analysis prioritized natural killer T cells and CD8+ and CD4+ T lymphocytes as relevant cellular contexts (Fig. 3d). These findings were interpreted at the trait level rather than assigned to individual loci without direct support. VariantAgent recovered 5 of 7 prespecified, directly comparable findings from the source study (71.4%), reproducing its principal genetic and cellular conclusions, although locus-specific experimental mechanisms were less completely recovered.

Using the same predefined path definition, blinded manual encoding identified 12 source-study paths, whereas VariantAgent assembled 608 paths linking variants to genes, molecular consequences, cellular contexts, perturbation evidence and pharmacological relationships. Of these, 566 contained only source-backed relationships, 18 combined source-backed and predictive evidence, and 24 included graph-nominated candidate links. VariantAgent recovered 3 of the 12 source-study paths; its total of 608 paths represented a net increase of 596 relative to the 12 paths encoded from the publication. This expansion reflects broader evidence integration rather than an equivalent number of experimentally validated mechanisms.

At rs9517712, fine-mapping, a whole-blood eQTL association and an AlphaGenome-predicted regulatory effect nominated TM9SF2 as an additional candidate gene (Fig. 3e). The absence of colocalization, direct regulatory interaction and perturbational support left this relationship unresolved. In contrast, rs34536443 supported a literature-backed path linking the protective T1D association to the TYK2 P1104A substitution and context-dependent attenuation of selected cytokine responses (Fig. 3f–h). Protein annotations and molecular-effect predictions complemented published functional evidence (Fig. 3f,g). CD8+ T-cell prioritization and the TYK2–deucravacitinib relationship remained separate contextual branches; the latter indicated target tractability rather than demonstrated clinical efficacy in T1D.

### VariantAgent reproduces published GWAS findings and expands downstream evidence

To evaluate generalizability beyond T1D, we systematically reanalysed 34 published GWAS^29–45^ spanning diverse diseases and quantitative traits (Fig. 4a). Source-study findings and VariantAgent outputs were compared across eight predefined evidence domains, including genetic signals, prioritized variants and genes, cellular contexts, mechanistic hypotheses, perturbation evidence and translational relationships. Recovery was assessed separately from evidence expansion so newly generated findings did not compensate for unrecovered source-study results. VariantAgent achieved a mean per-study recovery of 82.1% (95% CI, 75.3–88.9%; Fig. 4b).

**Figure 4.**
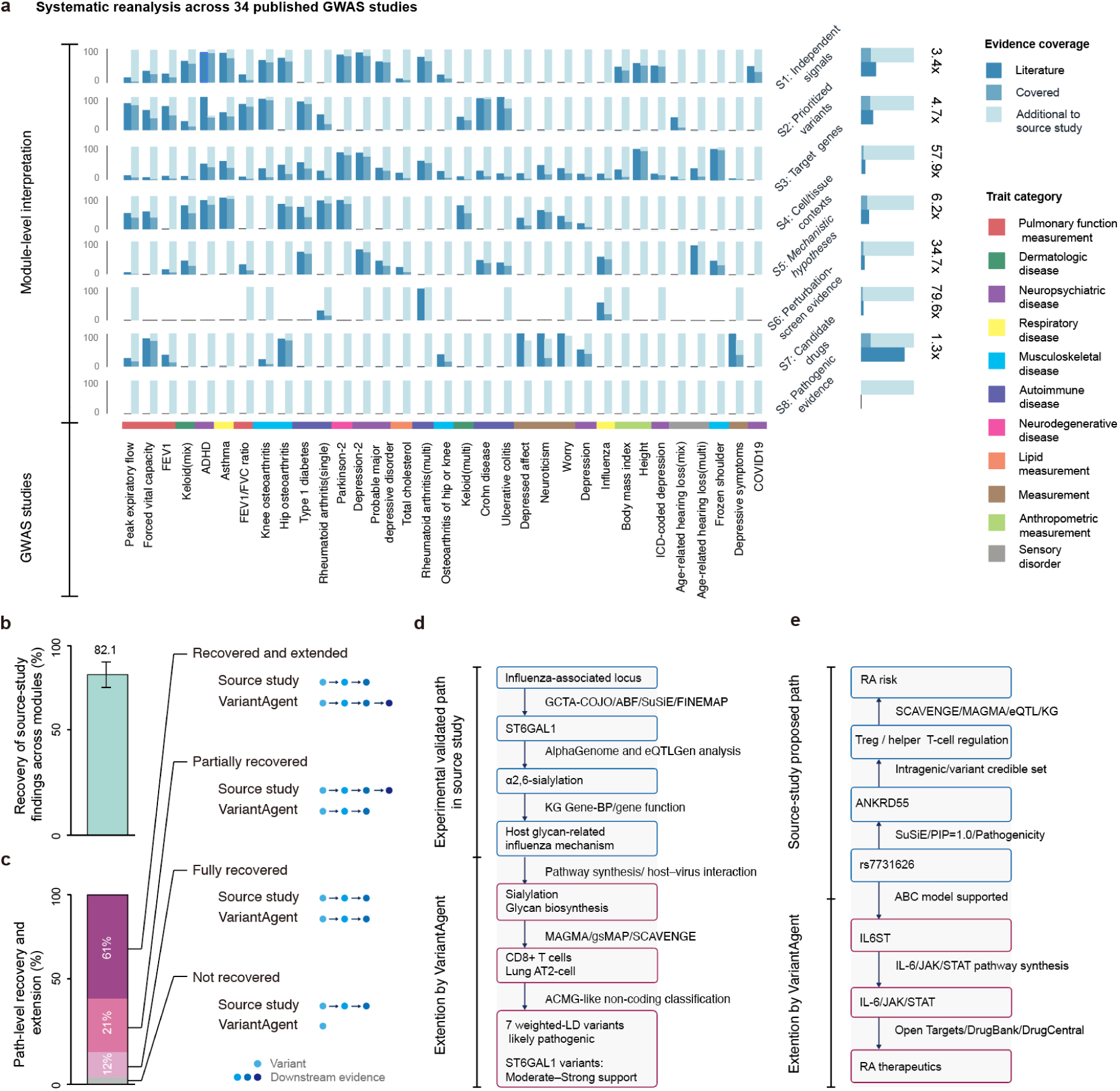
Matched reanalysis of published GWAS recovers reported findings and extends downstream biological interpretation. **a**, Systematic reanalysis of 34 published GWAS across eight post-GWAS evidence domains. Columns represent individual GWAS datasets and rows represent evidence domains. Within each study–domain combination, bars indicate evidence reported in the source study, evidence recovered by VariantAgent and evidence additional to the source study. The colour strip below the matrix denotes broad trait categories. The ratio of mean reanalysis to source-study evidence counts across eligible study–scenario comparisons is shown for each domain at right. **b**, Recovery of directly comparable source-study findings across evidence domains. The bar indicates the mean per-study recovery of 82.1%. **c**, Path-level recovery and extension of directly comparable source-study paths. Comparable paths were classified as not recovered, partially recovered, fully recovered, or recovered and extended; schematic paths illustrate the criteria used for each category. **d**, Representative recovered and extended locus-level path for influenza. The source study linked an influenza-associated locus to *ST6GAL1*, α2,6-sialylation and a host glycan-related influenza mechanism. VariantAgent computationally recovered the same locus–gene–mechanism axis using *de novo* fine-mapping, regulatory, expression and gene-function evidence, and extended the interpretation to broader sialylation and glycan-biosynthesis programs, disease-relevant CD8⁺ T-cell and lung AT2-cell contexts, and non-coding pathogenicity interpretation. **e**, Representative rheumatoid arthritis example showing recovery of a source-study path together with an alternative candidate branch from the same variant anchor. The source study linked rs7731626 to *ANKRD55*, Treg/helper T-cell regulation and rheumatoid arthritis risk. VariantAgent recovered this axis and additionally linked rs7731626 to *IL6ST* through ABC enhancer–gene mapping, followed by an *IL-6/JAK/STAT* pathway and therapeutic extension supported by pathway synthesis and drug-target resources. Blue outlines denote source-study and recovered relationships; magenta outlines denote evidence added by VariantAgent.

Uniform reanalysis revealed differences in evidence coverage across analytical domains (Fig. 4a). Across eligible study–scenario comparisons, the ratio of mean reanalysis to source-study evidence counts was 3.4-fold for independent signals and 4.7-fold for prioritized variants, but more markedly for prioritized genes (57.9-fold), tissue or cellular contexts (6.2-fold), mechanistic hypotheses (34.7-fold) and perturbation evidence (79.6-fold). Pharmacological links increased by 1.3-fold. These differences indicate broader downstream interrogation, not equivalent gains in biological validation.

At the level of variant-centred evidence paths, 739 source-study paths were directly comparable with automated outputs (Fig. 4c). VariantAgent fully recovered 155 paths (21.0%) and recovered and extended another 450 (60.9%), yielding an overall recovery-or-extension rate of 81.9%. Another 89 paths (12.0%) were partially recovered and 45 (6.1%) were not recovered. Paths identified exclusively through reanalysis were quantified separately and excluded from recovery calculations.

Representative examples illustrate reconstruction and expansion (Fig. 4d,e). For influenza, VariantAgent recovered the experimentally supported ST6GAL1–sialylation axis through independent genetic and functional evidence and extended its interpretation to glycan-related programs, cellular contexts and non-coding pathogenicity assessment (Fig. 4d). For rheumatoid arthritis, VariantAgent recovered the reported rs7731626–ANKRD55–T-cell relationship and identified an additional IL6ST–IL-6/JAK/STAT branch supported by enhancer–gene mapping, pathway annotations and drug-target evidence (Fig. 4e). These analyses show that systematic reanalysis can recover published findings while broadening the evidence available for interpretation.

### PGI makes accumulated post-GWAS evidence interrogable

Applying VariantAgent to 1,041 GWAS^46^ generated 530,949 standardized variant-centred evidence units and corresponding trait-level reports (Fig. 5a). PGI organized these outputs across 370 traits, preserving study context, analytical provenance and evidential status. Records derived from the same primary source were consolidated by provenance before cross-study comparison. The catalogue integrates genetic associations, variant-to-gene links, cellular contexts, molecular mechanisms, perturbation evidence and translational relationships.

**Figure 5.**
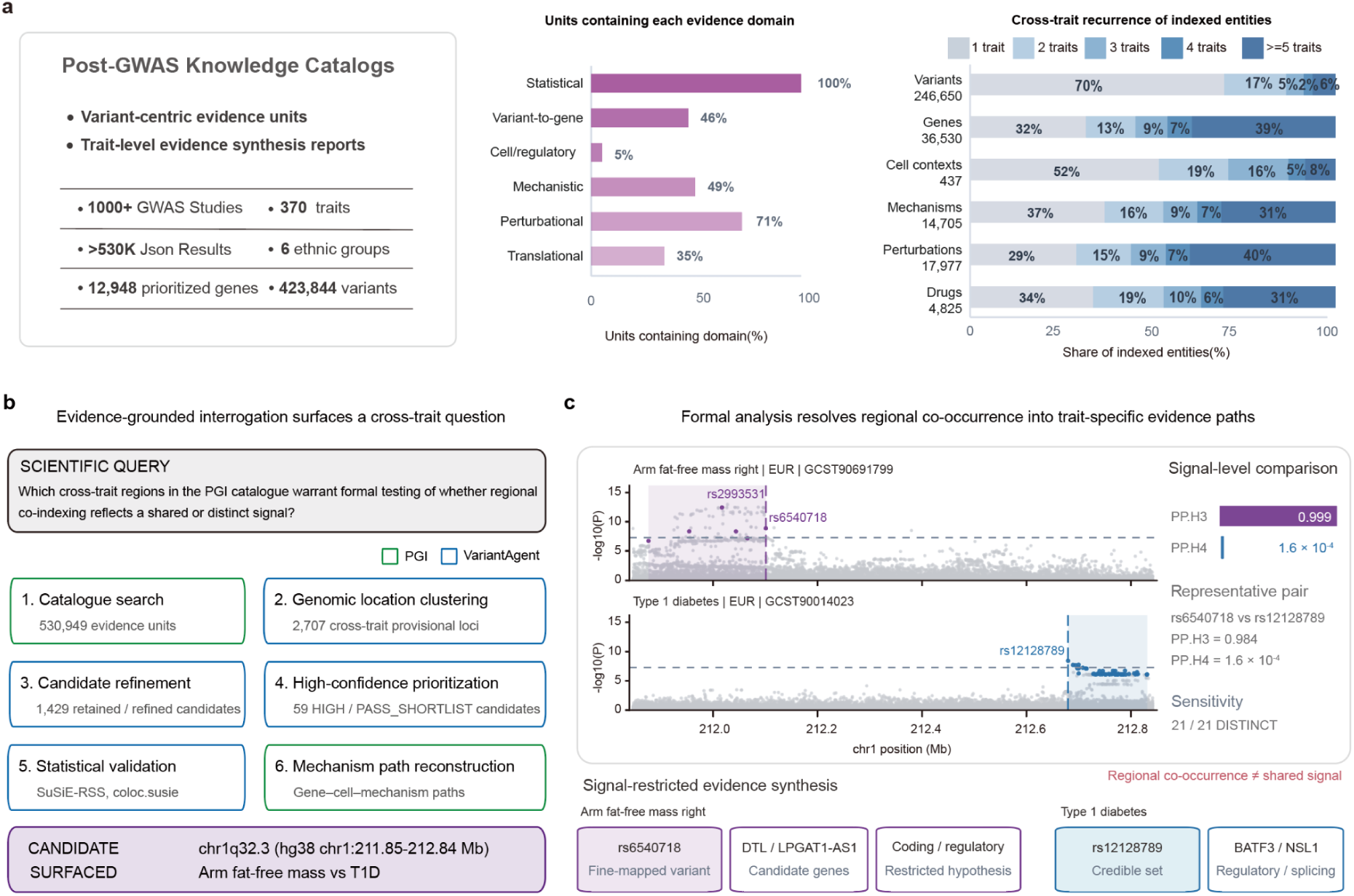
Accumulated post-GWAS evidence enables cross-trait interrogation and formal hypothesis testing. **a,** Scale and composition of the PGI post-GWAS knowledge catalogue. Variant-centred evidence units and trait-level evidence synthesis reports generated from 1,041 GWAS across 370 traits were indexed across statistical, variant-to-gene, cellular or regulatory, mechanistic, perturbational and translational evidence domains. Right, cross-trait recurrence of indexed variants, genes, cellular contexts, mechanisms, perturbations and drugs, grouped by the number of traits in which each entity was represented. **b,** PGI-unit-guided identification of a cross-trait hypothesis. Catalogue-wide screening of 530,949 evidence units identified 2,707 provisional cross-trait loci. Successive filtering for regional association support, study independence, fine-mapping evidence and downstream interpretability retained 1,429 candidates, including 59 high-confidence candidates for formal analysis. The chr1q32.3 region containing associations with right-arm fat-free mass and T1D was prioritized because regional co-occurrence was accompanied by discordant variant- and gene-level evidence. **c,** Formal statistical resolution and signal-restricted evidence synthesis at chr1q32.3. Analysis of the original regional GWAS summary statistics using SuSiE-RSS and signal-level coloc.susie supported distinct rather than shared association configurations for right-arm fat-free mass and T1D; all 21 sensitivity analyses favoured the distinct-signal model. Restricting PGI evidence to variants within the resolved configurations linked the body-composition signal to rs6540718 and *DTL/LPGAT1-AS1*, whereas the T1D configuration implicated rs12128789 and *BATF3/NSL1*. These relationships are shown as trait-specific candidate paths; cellular and direct mechanistic links that remained unresolved were not inferred.

An evidence-grounded conversational interface enables retrieval and synthesis of these records while retaining links to their analytical sources (Extended Data Fig. 5). For chronic obstructive pulmonary disease, iterative queries prioritized EIF3H while preserving uncertainty about the causal regulatory target. Cross-trait comparison of COVID-19 and influenza revealed limited overlap in prioritized variants and genes but convergence in CD8+ T-cell and alveolar epithelial contexts. Cross-ancestry comparison of Parkinson’s disease GWAS identified shared gene-level evidence involving SNCA and GBA alongside differences in locus and lead-variant representation. These examples illustrate evidence-grounded interrogation across loci, traits and populations without treating generated interpretations as primary evidence.

### Catalogue interrogation resolves an apparent cross-trait locus

We next investigated whether accumulated evidence could identify genetic questions requiring new statistical analysis. Querying the catalogue for regions represented across non-redundant traits but supported by distinct fine-mapped variants or downstream gene paths identified 2,707 provisional cross-trait candidates (Fig. 5b). Prespecified criteria for study independence, regional association, fine-mapping support and biological interpretability retained 1,429 candidates, of which 59 were prioritized for formal statistical evaluation. A candidate at chr1q32.3 contained associations for right-arm fat-free mass and T1D but suggested different fine-mapped variants and candidate-gene relationships.

VariantAgent reanalysed the original regional GWAS summary statistics using SuSiE-RSS and signal-level coloc.susie (Fig. 5c). Three right-arm fat-free-mass GWAS were compared with T1D across 21 sensitivity configurations. The body-composition associations resolved into two principal signals, whereas T1D showed a distinct configuration centred on rs12128789. All comparisons supported distinct rather than shared signals (PP.H3 ≥ 0.9989; PP.H4 ≤ 1.94 × 10⁻⁴).

Subsequent catalogue interrogation was restricted to variants within statistically resolved configurations. The body-composition signal included rs6540718 and candidate genes DTL and LPGAT1-AS1; the T1D signal implicated rs12128789 and correlated variants linked to BATF3 and NSL1, with regulatory and splicing-related support. Cellular contexts and direct molecular mechanisms remained unresolved. Thus, PGI identified an apparent cross-trait overlap that independent statistical analysis resolved into distinct signals and trait-specific biological hypotheses.

## DISCUSSION

This study demonstrates that heterogeneous post-GWAS analyses can be made systematic and comparable without requiring every study to follow the same analytical route. PGI couples domain-guided execution by VariantAgent to a common representation of the evidence produced. Across complementary benchmarks, matched reanalyses of 34 published GWAS and application to more than 1,000 datasets, VariantAgent reproduced expert-reported findings, broadened downstream evidence interrogation and generated standardized outputs that PGI organized across studies and traits. The central advance is the connection between conditional analytical execution and cumulative evidence production. Analytical routes can vary with the question and available data, while the resulting evidence remains traceable and reusable.

The benchmarks also expose an important distinction between answer accuracy and analytical validity. Comparator agents could reach plausible conclusions without executing the required method, preserving the intended variant set or correctly resolving genome build, allele orientation and ancestry-matched linkage disequilibrium. The separation between systems widened when a correct answer was required to be supported by a valid analytical trace. For computational claims, that trace forms part of the evidential basis for judging the conclusion. Scientific agents should therefore be evaluated not only by what they answer, but also by whether the analytical path to that answer is methodologically valid.

The matched reanalyses revealed that the largest differences in evidence coverage occurred downstream of association mapping. VariantAgent recovered a mean of 82.1% of directly comparable source-study findings. Uniform reanalysis produced relatively modest increases in independent signals and prioritized variants, but much larger increases in prioritized genes, mechanistic hypotheses and perturbation evidence. This asymmetry indicates that functional evidence remains far less consistently interrogated than genetic associations. PGI makes this broader evidence space accessible while preserving the maturity of individual relationships. In the T1D analysis, a literature-anchored coding path centred on TYK2 P1104A and an earlier-stage rs9517712-*TM9SF2* hypothesis could be represented within the same framework without assigning them equivalent evidential weight. Scale is thus achieved through systematic evidence interrogation rather than by treating additional candidates as established mechanisms.

The broader implication is that accumulated evidence can change the questions that become visible. Once interpretations from many GWAS are represented in a common, traceable form, cross-study discrepancies and recurrent relationships become objects of analysis in their own right. At chr1q32.3, conversational interrogation exposed an apparent cross-trait overlap that VariantAgent resolved against the original GWAS data into distinct signals linked to trait-specific biological hypotheses. PGI thereby extends the cumulative logic of human genetics beyond association mapping, allowing evidence generated by previous analyses to shape the questions asked next.

## METHODS

### VariantAgent architecture and workflow execution

VariantAgent comprises five agent types—Orchestrator, Planner, Executor, Reflector and Report Agent—organized into four Orchestrator-guided stages: planning, module execution and reflection, workflow-level reflection and revision, and reporting (Fig. 1b). The Orchestrator coordinates dependency-aware execution and review, while agents share inputs but retain separate execution contexts.

At workflow initiation, the Orchestrator launches the Planner, which constructs a structured execution plan based on the analytical objective and available data. The plan specifies the analytical modules to be performed, their required inputs and expected outputs, and the dependencies that determine their execution order. The Orchestrator interprets these dependencies during runtime to determine which modules are eligible for execution. Modules with satisfied dependencies can therefore be dispatched concurrently, whereas downstream modules remain blocked until the required upstream modules reach the appropriate terminal states. Each eligible module is assigned to a newly instantiated Executor, which performs the analysis and persists intermediate files, structured outputs and runtime logs as explicit workflow artifacts. After each module, an independent Reflector assessed observable artifacts for completeness, execution correctness and methodological validity, returning PASS, NEED_REVISION or SKIP_WITH_REASON. Revisions triggered a fresh Executor and review, with at most three execution–reflection cycles per module. A workflow-level Reflector then checked plan coverage, dependencies and cross-module consistency, requesting targeted re-execution when necessary. Following approval, the Report Agent synthesized validated artifacts and execution records into a traceable report.

More than 40 task-specific skills specified procedures, tools, outputs and quality criteria for Executors and Reflectors.

### VariantAgent model configuration and computational environment

VariantAgent used Claude Code v2.1.168 as its agent runtime and orchestration layer. The backend models evaluated in this study were Claude-Sonnet-4.6 and DeepSeek-V4-Pro, both of which support context windows of up to one million tokens. All agents in a workflow used the same backend model but distinct role-specific prompts, avoiding within-workflow model heterogeneity. Each dataset was analysed in a dedicated Docker container, preventing cross-dataset carryover of sessions, runtime states, processes and caches. Intermediate outputs were retained for provenance and downstream analysis.

### Post-GWAS analysis procedures

#### Scenario-oriented post-GWAS workflow

A scenario-oriented, variant-centred workflow linked GWAS signals to prioritized variants and genes, cellular contexts, mechanisms, experimental evidence and translational hypotheses. For each request, VariantAgent mapped the task to supported analytical scenarios using prespecified, scenario-specific applicability rules encoded in the domain SOPs. Scenarios ran only when inputs, resources and assumptions were sufficient; otherwise, applicability and execution states were recorded. Each run recorded the trait definition, source study, ancestry, sample size, genome build, input checksum, reference panels, software and database versions, analytical parameters and execution status. Intermediate outputs, quality-control results and provenance were retained to support traceability.

#### Input harmonization and quality control

The minimum fields required for data ingestion were variant identifier, chromosome, genomic position, alleles and P value. These fields supported variant normalization, locus annotation and evidence retrieval. Additional fields, including effect estimate, standard error, sample size and effect-allele frequency, were required for specific analyses such as meta-analysis, conditional signal detection and LD-based fine-mapping. Missing scenario-specific fields did not prevent ingestion of the dataset but rendered the corresponding analyses unavailable. Variants were standardized to chromosome-position-reference-alternative representation and linked to rsIDs where possible. Indels were left-aligned and minimally represented relative to the declared reference genome. Reference alleles were checked against a genome-build-matched reference sequence, and unresolved coordinate or allele conflicts were excluded from coordinate- and direction-sensitive analyses. Multiallelic variants were split into allele-specific records when unambiguous. Palindromic variants were retained for non-directional annotation but excluded from direction-sensitive analyses when strand orientation could not be resolved. Duplicate records were resolved using a prespecified hierarchy based on the author-designated primary analysis, completeness of sample-size metadata, variant-quality information and fixed source priority. Records with irreconcilable association statistics were flagged and excluded from fine-mapping rather than selected according to significance. Effect, non-effect, reference and alternative alleles were harmonized across GWAS, LD, QTL and functional resources. Directional molecular interpretations were made only when the GWAS effect allele could be aligned unambiguously to the corresponding molecular-assay allele. Otherwise, evidence was reported without directional claims.

#### GWAS meta-analysis and independent-signal detection

Meta-analysis was undertaken when at least two harmonizable cohort-level GWAS summary-statistic datasets were available. Cohorts were grouped by ancestry and compatible phenotype definition; genome-build and allele harmonization preceded pooling. Same-ancestry cohorts were combined using METAL standard-error weighting, with heterogeneity statistics retained. Ancestry-stratified results were preserved for downstream LD-dependent analyses. For multi-ancestry collections, aggregate results informed association summaries and locus identification, but LD-dependent conditional analysis and fine-mapping used ancestry-specific statistics and matched reference panels where available. LD reference panels were selected according to ancestry and genome build. An ancestry-matched 1000 Genomes reference panel or a documented study-specific panel was used by default. When no suitable panel was available, LD-dependent analyses were not performed or were reported only as sensitivity analyses with explicit limitations. Independent association signals were identified using GCTA-COJO for quality-controlled single-ancestry GWAS or same-ancestry meta-analysis results. Unless otherwise specified, genome-wide significance was defined as *P* < 5 × 10^−8^. Initial fine-mapping loci were defined around COJO or prespecified lead signals using fixed physical windows and lead-variant LD. Credible-set overlap was used only for *post hoc* locus consolidation and did not retrospectively redefine the initial analysis units.

#### Statistical fine-mapping

Single-ancestry summary statistics with compatible LD were analysed using complementary fine-mapping methods comprising approximate Bayes factors, SuSiE-RSS and FINEMAP. Unless otherwise specified, analyses used ±500-kb windows around independent or lead signals and generated 95% credible sets. Approximate Bayes factors were calculated using Wakefield’s approach. SuSiE-RSS and FINEMAP used harmonized summary statistics and ancestry-matched locus-specific LD matrices. Analyses with insufficient variant overlap, unresolved allele mismatches, incompatible LD matrices or invalid sample-size information were assigned partially_successful, failed or not_available status according to whether interpretable quality-controlled outputs remained. Prespecified fallback procedures were used only after resolving input and LD quality-control failures; both initial and fallback results were recorded.

Formal trans-ancestry fine-mapping was performed with SuSiEx when ancestry-stratified GWAS summary statistics and corresponding ancestry-specific LD matrices were available. SuSiEx jointly modelled association evidence and LD patterns across ancestries within harmonized locus boundaries. Analyses based only on mixed-ancestry aggregate summary statistics were not treated as formal trans-ancestry fine-mapping. ABF outputs were reported as SNP posterior probabilities, whereas SuSiE and FINEMAP outputs were reported as posterior inclusion probabilities. A variant was considered method-supported when its method-specific posterior probability or PIP was >0.1. Variants supported by at least two of ABF, SuSiE and FINEMAP were termed multi-method-supported fine-mapped candidates. Variants prioritized by formal ancestry-aware methods were reported separately as trans-ancestry-prioritized candidates.

#### Variant-to-gene prioritization

Candidate genes were prioritized by integrating coding and splice consequences, regulatory links, MAGMA gene-level association, molecular-QTL annotations, chromatin interactions, expression in trait-relevant tissues, perturbation evidence, curated evidence and knowledge-graph relationships. V2G evidence was organized using a prespecified qualitative hierarchy. Direct protein-altering or splice-disrupting consequences within a candidate gene represented the most direct evidence. Fine-mapped non-coding variants linked to genes through enhancer-promoter, activity-by-contact or chromatin-interaction evidence in a relevant tissue or cell type formed the next tier. MAGMA association, allele-aligned molecular-QTL annotations, concordant expression or chromatin evidence and non-colocalized QTL overlap were treated as supportive evidence. Nearest-gene assignments, weak regulatory overlaps and knowledge-graph-only paths were considered exploratory. Multiple candidate genes were retained when the available evidence supported competing interpretations of the same locus. Molecular-QTL overlap was treated as annotative V2G evidence and was not described as colocalization. Where formal colocalization results were available from a task-specific analysis or an external source, they were recorded separately and required compatible GWAS and QTL summary statistics, shared variants, appropriate LD information, prespecified priors and posterior thresholds. Formal colocalization was not systematically performed across all analyses and was therefore not used as a universal highest-tier criterion. Evidence records retained both modality (statistical, regulatory, coding, molecular-QTL, experimental or computational) and origin (direct analysis, curated resource, literature, user input or graph inference).

#### Cellular context and pathway analysis

Cell- and tissue-context analyses were performed when suitable trait-relevant datasets and required statistical inputs were available. Gene-set and cell-type enrichment analyses were performed with MAGMA, with the background gene set, marker source and multiple-testing correction recorded. Single-cell ATAC-seq localization was performed with SCAVENGE by integrating fine-mapped variant posterior probabilities with a genome-build-matched single-cell ATAC-seq peak matrix and corresponding cell annotations. Spatially resolved mapping of trait-associated cells was performed with gsMAP by integrating GWAS summary statistics with trait-relevant spatial transcriptomic data. Analyses were restricted to spatial datasets with appropriate tissue context, sample provenance and cell or regional annotations. Cell, tissue and spatial enrichment provided contextual rather than locus-specific mechanistic evidence. Such findings were not assigned to individual variants without independent locus-level support.

#### Non-coding mechanism annotation

Prioritized non-coding variants with unambiguous genomic coordinates and allele definitions were evaluated for regulatory and splicing mechanisms. AlphaGenome was used to predict potential variant effects on regulatory molecular phenotypes, including chromatin accessibility, transcription-factor binding, gene expression and other sequence-dependent regulatory features, whereas SpliceAI was used to assess splice-altering potential. Model outputs were interpreted together with regulatory-element and chromatin-state annotations, enhancer-promoter links, activity-by-contact evidence, chromatin interactions and molecular-QTL evidence. Non-coding mechanisms were proposed only when variant identity, allele orientation, candidate-gene linkage and cellular context were consistent; predictions alone were not treated as experimentally established effects.

#### Protein functional impact annotation

Eligible coding variants were annotated with VEP or SnpEff and mapped to transcripts and protein products. Predicted loss-of-function and splice-disrupting consequences were retained as consequence-level evidence; ineligible variants were marked not_applicable rather than negative. ProtVar was queried for missense substitutions with valid UniProt accession, residue position and amino-acid change, providing residue-level functional annotations. Protein evidence distinguished sequence consequence, deleteriousness, residue function, structural predictions and experimentally supported mechanisms; computational scores were not treated as functional validation.

#### Experimental, clinical and translational evidence

Experimental support was retrieved from variant-level reporter assays, massively parallel reporter assay (MPRA), multiplexed assays of variant effect (MAVE), saturation-mutagenesis datasets and gene-level CRISPR screens. Retrieved evidence was distinguished from lack of resource coverage: failure to identify a matching record in a queried resource was treated as absence of retrieved support and not as experimental evidence against the candidate. Clinical classification was restricted to clinically oriented analyses with appropriate variant-level evidence; ACMG/AMP- or ClinGen-informed categories were not routinely assigned to common GWAS variants. Drug and translational annotations distinguished direct drug–target relationships from pathway associations, pharmacogenomic evidence and therapeutic hypotheses; target tractability was not equated with disease-specific efficacy.

#### Workflow reporting and analytical status

Each run reported module applicability, execution status, quality control, artifacts and limitations. Only quality-controlled, provenance-verified outputs supported findings; failed, skipped or unavailable analyses did not count as evidence. Reports linked findings to outputs, tools, parameters and resource versions. Evidence-unit construction and cross-variant synthesis occurred during catalogue generation; computational nominations were not treated as validation.

### Construction and use of the biomedical knowledge graph

Biomedical knowledge graph construction. We constructed GWAS-KG in Neo4j to connect genetic variants, genes, phenotypes, biological functions, drugs, anatomical structures and publications through source-backed relationships. The graph comprises 8,935,910 nodes across 12 entity types and 28,394,957 edges across 47 relationship types. Of these, 8,709,690 are variant nodes.

GWAS-KG integrated the GWAS Catalog, ClinVar, Open Targets, PhyloP, CADD, gnomAD and PrimeKG, covering variant–trait associations, clinical significance, variant-to-gene and drug-target links, conservation, functional impact, population frequencies, gene constraint and broader biomedical relationships. Provenance was retained after normalization.

Entity normalization. Variants were normalized to rsIDs and GRCh38 chromosome, position and alleles; genes to HGNC symbols with Ensembl and NCBI cross-references; and phenotypes to MONDO with EFO mappings. Other entities used DrugBank/ChEMBL (drugs), GO (functions), Reactome (pathways), UBERON (anatomy), HPO (indications), PubMed IDs (publications) and MeSH (exposures). Duplicate canonical entities were merged while retaining provenance; ambiguous mappings remained separate.

Relationship construction. Source-specific typed edges captured genetic associations, variant–gene links, functional annotations, tissue expression, ontology relationships and pharmacological interactions, retaining available scores, P values, evidence counts and provenance. Source-backed and inferred links remained distinct.

Knowledge graph query. GWAS-KG was queried using Cypher in Neo4j Community Edition (Kernel v2026.03.1). A Model Context Protocol interface exposed graph queries and algorithms to VariantAgent while retaining source provenance.

#### Graph-based analysis

Neo4j local subgraphs were projected around candidate variants, genes and target phenotypes, retaining direct and multi-hop relationships among variants, genes, phenotypes, drugs, pathways and GO terms. Neo4j Graph Data Science provided PageRank, Personalized PageRank and Node2Vec. PageRank measured overall network importance:

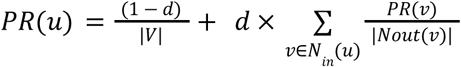

where *d* denotes the damping factor and was set to 0.85.

Personalized PageRank was used to estimate the network proximity between candidate genes and target phenotypes:

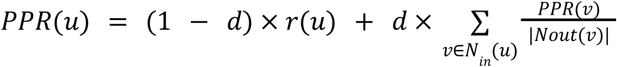

where,

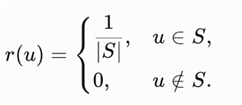

when node *u* belongs to the phenotype seed set *S*, and *r*(*u*) = 0 otherwise. All target phenotype nodes were assigned equal restart probabilities.

Node2Vec used 32-dimensional embeddings, walk length 10, 20 walks per node and p = q = 1 for topological similarity and multi-hop analysis.

#### Graph-based gene prioritization

A composite score combined phenotype-specific network proximity, phenotype–gene association strength, supporting evidence counts and gene loss-of-function constraint:

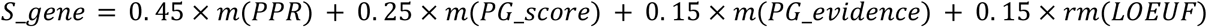

Forward Min–Max normalization was applied to Personalized PageRank, phenotype–gene association strength and supporting evidence counts; reverse Min–Max normalization was applied to loss-of-function observed/expected upper bound fraction (LOEUF), for which lower values indicate stronger constraint.

Weights were selected empirically by comparing candidate schemes, not by supervised fitting: 0.45 for phenotype-specific Personalized PageRank, 0.25 for phenotype–gene association strength, 0.15 for supporting evidence counts and 0.15 for LOEUF-based constraint. Genes were ranked by the resulting composite score.

### Generation of post-GWAS knowledge catalogues

Post-GWAS catalogues comprised full-process reports (Markdown), variant-centred evidence units (JSON) and trait-level synthesis reports (Markdown), retaining identifiers, provenance, resource versions, status and evidence type.

Full-process analysis reports. Versioned reports recorded study metadata, parameters, quality control, source-linked conclusions and reasons for incomplete analyses; unvalidated outputs were not counted as evidence.

Variant-centred evidence units. Each unit was keyed by rsid, trait and source_GWAS, with harmonized coordinates, alleles and genome build. Records covered associations, independent signals, fine-mapping, V2G links, mechanisms, pathways and experimental or translational evidence. V2G support distinguished coding/splicing, MAGMA, QTL, enhancer, chromatin and graph sources; mechanism evidence distinguished regulatory, splicing and protein effects. Pathway and trait-level cell enrichment were not attributed to individual variants without direct support. Records retained source, version, tool, parameters, evidence class and status.

#### Trait-level evidence synthesis reports

Trait-level reports were generated from collections of variant-centred units, grouping supported relationships and unresolved hypotheses without introducing unsupported evidence. Candidate variant-gene-mechanism relationships were ranked using a prespecified rule-based framework that considered genetic support, the directness of V2G assignment, convergence across independent evidence sources and analytical scenarios, molecular-mechanism support and unresolved conflicts. Evidence records derived from the same original source were counted once, missing evidence was treated as reduced coverage rather than contradictory evidence, and conflicting evidence was retained as an explicit ranking attribute. This scientific candidate ranking was performed independently of the query-relevance scores used for catalog retrieval.

External checks queried PubMed, the GWAS Catalog and Open Targets for matching variant–gene–trait relationships, retaining source identifiers and query provenance. Candidates were classified as known mechanisms, known associations, database-unreported candidates or unresolved. Database-unreported denoted absence from the queried sources at the recorded date, not proven novelty; incomplete searches remained unresolved. Reports presented ranked variant–gene–mechanism relationships, supporting evidence, known relationships and candidates requiring follow-up.

Catalogue indexing and evidence retrieval. Units were indexed by rsID, trait and source_GWAS; exact matches preceded contextual retrieval, with provenance-preserving ranking and deduplication.

### Full-process evaluation of agent performance

Full-process scoring assessed observable plans, commands, tool calls, parameters, intermediate outputs, failures and reports using a predefined rubric evaluated by Codex with GPT-5.5. Unsupported claims of execution were not credited. The rubric comprised two aspects and eight dimensions. Execution capability included Resource Awareness and Execution Planning, Data Preprocessing and Cross-Resource Harmonization, Software and Tool Execution Performance, and Resource Coverage and Diversity. Execution rationality included Statistical Rigor and Appropriateness, Variant-to-Gene Multi-Source Endorsement, Molecular Mechanism Resolution, and Integrated Evidence Report and Hypothesis Generation. Each dimension was scored independently on a 0-100 scale. The two aspect scores were calculated as the mean of their four component dimensions, and the overall score was calculated as the mean across all eight dimensions. Scoring covered appropriate tool selection and execution, genome-build and allele harmonization, ancestry and resource compatibility, statistical validity, multi-source gene and mechanism evidence, and traceability. Tool states distinguished success, partial success, failure, justified or unjustified skipping, and unavailability; planned but unexecuted work did not count. We also reported tool and resource usage, execution failures, scenario coverage and evidence traceability. These scores assessed adherence to the rubric, not experimental validation of biological conclusions.

Three blinded experts independently scored outputs from five GWAS tasks across eight dimensions. Expert consensus was the mean score; inter-rater reliability and agreement with agent scoring were assessed using a two-way random-effects absolute-agreement ICC, with ICC(A,k) as the primary agreement measure.

### Published genetic-reasoning benchmarks

We compiled 313 tests from published GWAS-related question-answering datasets associated with Biomni, GenomeArena and SDE. The benchmark comprised 230 phenotype-gene, 43 variant-phenotype and 40 variant-gene questions. Records were standardized by question, reference answer, relation type and source dataset; duplicates were removed, answer formats and equivalent labels harmonized, and phenotypes, genes and variants normalized to ontology or canonical identifiers where possible. GWAS-KG and supporting resources retrieved and ranked relationships without access to reference answers. Accuracy was scored consistently across systems, overall and by relationship type; the task assessed relationship retrieval, not full post-GWAS execution.

### GWAS-centric benchmark

We constructed a GWAS-centric benchmark from ten GWAS summary-statistics datasets distinct from those used in the full-process evaluation. Datasets were selected for availability, sample size, phenotype clarity and diversity, and checked for required fields and analytical compatibility. Candidate questions and multiple-choice options (A–D) were drafted with LLM assistance using study-specific information and BixBench-inspired examples. Manual review resolved ambiguity and duplication and checked alignment, task coverage and difficulty. Reference answers were curated from source publications and checked against the supplied GWAS datasets and, where applicable, expected analytical results. The final benchmark comprised 26 questions across ten GWAS datasets, covering conditional association, fine-mapping, gene and pathway prioritization, cellular context, sequence and protein function, drug evidence and pathogenicity. Agents had to interpret study-specific inputs, execute appropriate analyses when needed, harmonize evidence and produce free-text answers.

Each system received the benchmark question together with the corresponding GWAS summary statistics, and each system-question pair was evaluated in five independent runs. All systems generated free-text responses. Final responses and available traces (analytical steps, tool calls and intermediate outputs) were retained. GPT-5.5 assessed answers using three LLM-as-a-judge protocols; process measures used execution traces. For open answers, GPT-5.5 assigned binary correctness against curated references without viewing traces. For two multiple-choice protocols, each question had one correct option and three manually reviewed plausible distractors; the judge received the response and execution trace. Forced choice required selection of A–D; the refusal-permitted protocol also allowed E (insufficient information/unable to determine) when none of A–D was supported. A run was correct only when the selected option matched the reference answer.

Codex with GPT-5.5 scored complete traces against a prespecified process rubric independently of answer correctness. Five process metrics captured target-method execution, harmonization, evidence-chain completion, fallback recovery and process-validated accuracy, separating correct answers from those supported by appropriate analysis. Three experts independently assigned binary correctness labels; majority vote defined the expert reference, and agreement with the agent-derived binary label was calculated across items.

### Comparator systems and benchmark execution

Comparator systems were evaluated at two levels. For the 313-question published genetic-reasoning benchmark, five models—DeepSeek-V4-Pro, Qwen-3.7-Max, Kimi-2.6, Claude-Sonnet-4.6 and GPT-5.4—answered directly without agents or external tools. Considering accuracy, API token prices and per-question token use, we selected Claude Code + DeepSeek-V4-Pro + skill library for subsequent full-process and GWAS-centric analyses. Agentic comparators included Claude Code + DeepSeek-V4-Pro, OpenCode + DeepSeek-V4-Pro, Claude Code + Tool Universe + DeepSeek-V4-Pro, Biomni v0.0.8 + DeepSeek-V4-Pro, and Biomni + Claude-Sonnet-4.6 (included to reflect Biomni’s published Claude-family setting). The complete 313-question comparator set also included direct answers from all five models and Claude Code or OpenCode paired with the skill library and each of the five models. For the full-process post-GWAS and GWAS-centric evaluations, VariantAgent was compared with Claude Code (general-purpose coding agent) and Biomni (biomedical tool-augmented agent). All three used DeepSeek-V4-Pro as the sole base model. Differences in native tools and accessible resources were retained as components of each system’s capability rather than artificially equalized.

Systems received identical questions, data and instructions. Full-process and GWAS-centric inputs comprised supplied GWAS summary statistics, with ancestry, genome build and phenotype specified. Systems could write only within their working directories and access no other local files. Network access was allowed for required methods and resources (for example, 1000 Genomes LD panels), but prohibited for retrieving answers, source papers, abstracts, supplementary materials, external-model conclusions or search-engine results. Answers derived from each system’s analysis. Budgets and timeouts were recorded, with no post-initialization human intervention and sufficient time for long-running tools. For the broader evaluation, a prespecified subset of tasks was rerun five times in independent sessions to assess stability, while the remaining tasks were evaluated once; the 26-question GWAS-centric benchmark was separately evaluated with five independent runs per system–question pair. Runs did not share intermediate artifacts or traces. Comparisons evaluated end-to-end agent systems, whose base models, tool ecosystems and resources were not identical; results should not be interpreted as controlled comparisons of foundation models alone.

### Full-process post-GWAS analysis and evidence integration for T1D

T1D GWAS summary statistics came from a European-ancestry study of 18,942 cases and 501,638 controls. Inputs were harmonized to GRCh38 coordinates, alleles and rsIDs; COJO and fine-mapping used effective N = 61,427. Ten scenarios covered fine-mapping, V2G, cell context, sequence function, perturbation, drug evidence, ACMG-style interpretation, graph analysis, external validation and reporting, with module-level QC. Independent signals were identified using GCTA-COJO and the 1000 Genomes EUR GRCh38 LD panel (--cojo-slct, --cojo-p 5e-8, --cojo-wind 10000). The analysis identified 324 COJO-independent signals, of which 125 met P < 5 × 10−8 in both marginal and conditional analyses.

Fine-mapping used ABF (coloc::finemap.abf, prior p1 = 1 × 10−4, ±500 kb), SuSiE-RSS (L = 10, coverage = 0.95, estimate_residual_variance = TRUE, max_iter = 500, min_abs_corr = 0.5) and FINEMAP v1.4.1 (stochastic search, 1000 Genomes EUR locus-specific LD, ±500 kb, up to ten causal variants). SuSiE converged for 236 of 324 loci. A posterior probability or PIP threshold of 0.1 yielded 910 variants supported by at least two methods, including 52 supported by all three; REF/ALT validation passed for all 910. V2G prioritization integrated proximity, MAGMA, eQTLGen, eQTL Catalogue, ABC enhancer–gene links, pQTL evidence and graph relationships. ABC enhancer maps spanned 131 cell types; gene-level support was retained by source rather than inferred from overlap counts alone.

Gene-level functional interpretation used Enrichr over-representation analysis with GO, KEGG, Reactome and WikiPathways. Cell-context analyses combined MAGMA enrichment of 1,893 MSigDB C8 gene sets, gsMAP on Visium HD pancreas data (11,025 spots), and SCAVENGE on pancreatic snATAC-seq (GSE184462; 7,319 cells, 38 cell types) with blood validation (GSE239799; 5,977 cells). MAGMA enrichment was not significant after Bonferroni correction; SCAVENGE used the top 1% of g-chromVAR cells as seeds per fine-mapping method.

Coding and non-coding mechanisms were evaluated for 910 high-confidence variants using AlphaGenome (11 epigenomic modalities), SpliceAI (ds_max ≥ 0.20 threshold), VEP, SnpEff and Variant2Protein for eligible missense variants. Perturbation evidence for these variants and the top 50 genes was queried in BioGRID ORCS, MaveDB, MPRAvarDB, satMutMPRA and MPRAcollect; absence of matched records was distinguished from negative functional evidence.

Drug evidence for the top 50 genes was queried in Open Targets, ChEMBL, ClinPGx/PharmGKB and DrugCentral, with limited coverage for many MHC/HLA genes. ACMG/AMP 2015 criteria were applied to ten coding missense variants; an Ellingford 2022-style framework adapted for common GWAS alleles was applied to 896 non-coding variants. Allele-frequency evidence was considered explicitly in interpretation.

Knowledge-graph analysis used the 910 high-confidence variants and T1D phenotype MONDO:0007454/EFO:0009756 to prioritize 50 candidate genes. Graph-derived pathway and drug links were treated as contextual hypotheses, not causal evidence. Variant-centred evidence units retained gene-supported, functional-rescue and exploratory candidate categories; 66 candidates were checked against GWAS, PubMed and Open Targets sources. Evidence chains preserved provenance and distinguished direct functional support from QTL and computational hypotheses.

### Systematic reanalysis of published GWAS

#### Selection and standardized reanalysis of published GWAS

We reanalysed 34 published GWAS across respiratory, dermatological, neuropsychiatric, musculoskeletal, autoimmune, inflammatory, neurodegenerative, lipid, anthropometric and sensory traits. Selection required available summary statistics, diverse phenotypes, adequate sample sizes and metadata, and sufficiently detailed published post-GWAS results for comparison. Source findings were manually curated from articles and supplements but withheld during standardized VariantAgent reanalysis. To reduce direct source-study leakage, the source publication and supplementary materials were excluded from the analysis context, and records explicitly attributable to the evaluated study through PMID, GWAS Catalog accession (GCST) or other study-specific identifiers were masked where such provenance information was available. VariantAgent retained public evidence retrieval, so indirect source-study leakage through secondary resources could not be excluded; agreement therefore indicated recovery under controlled retrieval, not fully blinded independent replication.

Results were organized into eight analytical categories: independent association signals, fine-mapped candidate variants, prioritized candidate genes, cell or tissue contexts, *in silico* mechanisms, perturbation-screen evidence, candidate drugs and pathogenicity-related evidence. Source and reanalysis records were compared by category after within-study deduplication. Findings shared by source and reanalysis were recovered; reanalysis-only findings were additional; source-only findings were not recovered. These classifications describe correspondence and expansion of analytical coverage; recovered does not necessarily imply an independently rediscovered evidence source, and additional evidence does not by itself imply biological novelty or validation.

#### Scenario-level comparison and calculation of evidence expansion

Scenario-level analytical coverage was assessed by comparing entity or evidence-record counts across the 34 matched reanalyses. Within each study–scenario pair, source and reanalysis outputs were normalized to comparable units (signals, variants, genes, cellular contexts, mechanisms, perturbations, drugs or pathogenicity records) and deduplicated. Study-scenario combinations were considered eligible for ratio-based comparison when the source publication provided a non-zero countable output and the source-study and reanalysis outputs were sufficiently comparable. Counts had to represent the same output class, although evidence sources or prioritization methods could differ. Study-scenario combinations with zero source-study counts but non-zero reanalysis counts were retained as qualitative expansions but excluded from ratio calculations. Narrative-only, uncountable or non-comparable source outputs were also excluded. No pseudo-counts or continuity corrections were applied. For each scenario, source-study and reanalysis counts were restricted to eligible study–scenario combinations. Scenario-level expansion was then calculated as the ratio of the mean reanalysis count to the mean source-study count across the same eligible studies. With identical eligible studies, this equals the ratio of summed counts. Ratios measured analytical breadth, not biological validity.

#### Matching of variant-centred evidence relationships

Variant-centred paths were extracted separately from source publications and VariantAgent outputs, using the focal variant, downstream entities and relationship themes. Rule-based matching compared structured paths by shared focal variants, downstream entities and relationship themes, not unrestricted document text. At the broad comparison level, relationships were assigned to four categories: source-study variants not recovered in the reanalysis; variants identified only in the reanalysis; shared variants linked through downstream paths not represented in the source study; and shared variants for which source-study paths were reproduced and extended in the reanalysis. At the path level, relationships were classified as partially reproduced (shared focal variant and some downstream structure), fully reproduced (shared variant and principal downstream relationship), extended (reproduced core path plus additional entities or evidence), or previously unreported (no explicit source-study counterpart). Categories reflected agreement and expansion rather than exact path order; classifications were manually reviewed.

### Conversational interrogation of the post-GWAS knowledge catalogues

#### Catalogue composition and ingestion

PGI indexed 530,949 variant-centred evidence units from 1,041 GWAS, together with full-process and trait-level reports, for provenance-aware conversational retrieval. Retrieval and ranking. Natural-language queries were parsed into variants, traits, source studies, evidence categories and free-text entities. Exact identifier and full-text matching were complemented by semantic retrieval to identify relevant evidence fragments while preserving study identity.

Each evidence unit was uniquely identified by variant, trait and source study. Retrieval kept study- and ancestry-specific units separate. Fragments were ranked by entity match, trait consistency and relevance, then deduplicated with bounded retrieval per unit. Method, version, reference panel and parameter provenance accompanied the returned evidence. Responses cited retrieved fragments and flagged conflicting or insufficient evidence without inferring unsupported links.

#### Evidence-grounded response generation

Each response retained a retrieval-derived trace of the reports, evidence units and analytical modules actually supplied as context, rather than retrospectively generated citations. We illustrated iterative single-trait, cross-trait and cross-ancestry queries (Extended Data Fig. 5). The interface distinguished evidence types from hypotheses without substituting for statistical or experimental validation.

### PGI-driven cross-trait hypothesis discovery and statistical resolution

Variant-centred evidence JSON files were indexed by GRCh38 coordinates where available, trait, GWAS source, fine-mapping, V2G, cell-context, mechanism and provenance fields; summary-only files were excluded. Variants were clustered by genomic position, merging adjacent variants within 250 kb up to a maximum 1-Mb locus span. The MHC region was excluded. Loci represented by at least two normalized, non-redundant trait labels were retained as provisional cross-trait loci.

Within each provisional locus, we summarized evidence by trait and recomputed fine-mapping support from unit-level ABF posterior probabilities, SuSiE and FINEMAP PIPs, and credible-set membership. Trait pairs were compared by variant sets, top variants, candidate genes, V2G grades, and cell and mechanism annotations. Pairs with non-overlapping fine-mapped variants and candidate genes, low weighted variant overlap, and interpretable downstream differences formed the strict cross-trait shortlist. Shortlists were reassessed for regional association, significant-variant counts, fine-mapping, V2G, cell and mechanism evidence, study independence and data availability. We excluded near-duplicate traits, pairs lacking regional association in either trait, weak fine-mapping or interpretable V2G evidence, and likely overlapping study samples. Remaining pairs were labeled PASS_SHORTLIST or RETAIN_FOR_REVIEW. High-confidence candidates came from PASS_SHORTLIST after collapsing redundant pairs to one representative per provisional locus.

For chr1q32.3, we extracted original regional GWAS summary statistics for right-arm fat-free mass and T1D at PL01_211846683_212841862, harmonized coordinates to GRCh38, matched alleles across traits and intersected variants with the 1000 Genomes EUR LD reference. Each trait was independently fine-mapped using SuSiE-RSS; coloc.susie then compared all inferred signals between traits. Statistical classification used only original summary statistics, LD and SuSiE/coloc results, not downstream PGI annotations. Sensitivity analyses varied region boundaries, LD-variant inclusion and SuSiE settings. Distinct signals required consistently high H3 and low H4 support across valid configurations. Interpretation followed statistical classification, was limited to resolved variants and distinguished contextual annotations from direct mechanistic evidence.

## DATA AVAILABILITY

Metadata and source identifiers for the 1,041 catalogue-scale GWAS are available through the PGI portal at pgi.aigenomicsyulab.com. The PGI variant-centred evidence units generated in this study are also available through the PGI portal.

## CODE AVAILABILITY

The source code for VariantAgent is available from the corresponding authors upon reasonable request. PGI is accessible at https://pgi.aigenomicsyulab.com/.

## AUTHOR CONTRIBUTIONS

F.Y. conceived and designed the study. L.Q., S.G., X. Wang, and F.Y. designed and developed the agent framework and post-GWAS skills. L.Q., S.G., Y.Y., Y.H., and X.L. conducted data analyses, benchmarking, and hypothesis validation. Y.Y. and Y.H. developed the PGI web platform and retrieval algorithm, and X.L. developed the GWAS-KG. X. Wan, X.Z., Q.Z., T.H., Y.Z., F.C., and J.H. curated the post-GWAS analytical workflows. L.B., E.L., P.Y., J.W., Z.T. and L.W. provided constructive advice on the study design and agent framework. F.Y., L.Q., X. Wang, Y.Y., S.G., X.L., and Y.H. wrote the manuscript with input from all authors. F.Y., L.B., X. Wang, and E.L. supervised the study.

## ADDITIONAL INFORMATION

Correspondence and requests for materials should be addressed to Fulong Yu.

## ACKNOWLEDGEMENTS

We are grateful to members of the Yu laboratory and numerous colleagues for valuable comments and suggestions. This work was supported in part by grants GZNL2024A01003 and GZNL2023A02002 from the Major Project of Guangzhou National Laboratory, grant 2024B1515020080 from Guangdong Basic and Applied Basic Research Foundation, grant 32470634 from the National Natural Science Foundation of China, grant KY012023362 from the Talent Research Funding Project of Guangdong Provincial People’s Hospital. We acknowledge the Data Science Platform of Guangzhou National Laboratory and the Bio-medical Big Data Operating System (Bio-OS) for technical support and for providing access to the computational resources essential to this study.

## Extended data

**Extended Data Fig. 1.**
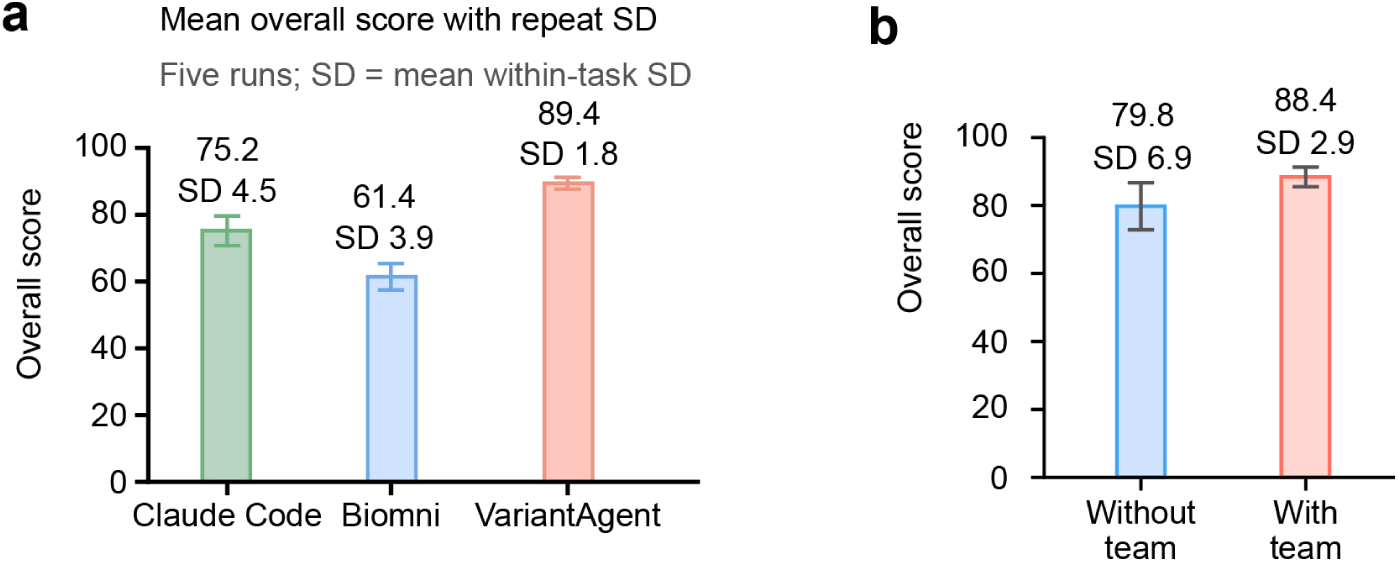
Run-to-run stability and contribution of the agent-team architecture to full-process performance. **a,** Repeated full-process evaluation across three representative GWAS datasets, each analysed independently five times. Bars show the mean overall score across repeated runs; error bars indicate the mean within-task standard deviation. VariantAgent achieved the highest mean score and the lowest run-to-run variability compared with Claude Code and Biomni. **b,** Ablation of the agent-team architecture under matched execution conditions. Bars show the mean overall score with and without the coordinated agent team; error bars indicate standard deviation across repeated runs. Removing the team architecture reduced overall performance and increased variability, supporting a contribution of coordinated planning, execution and review beyond access to the same analytical resources.

**Extended Data Fig. 2.**
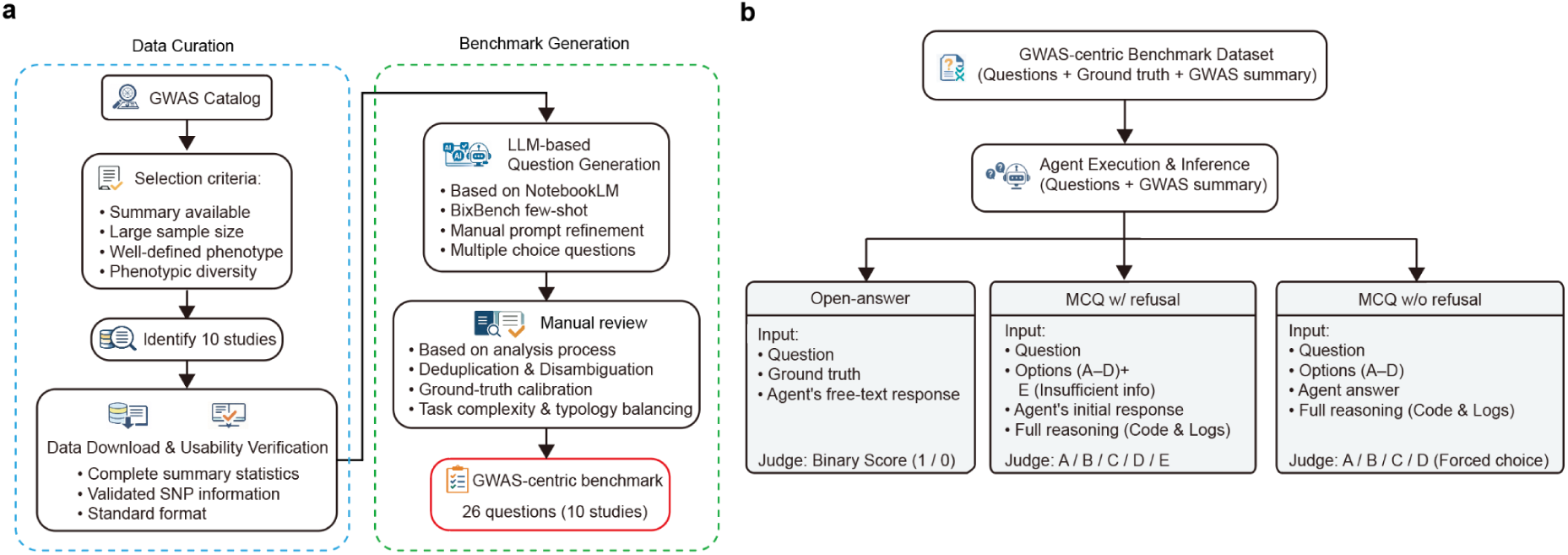
Construction and evaluation of the GWAS-centric benchmark. **a**, Curation and generation of the GWAS-centric benchmark. An independent set of ten GWAS summary-statistic datasets was selected on the basis of summary-statistic availability, sample size, phenotype definition and phenotypic diversity. Downloaded summary statistics were further assessed for completeness, variant information and format usability. Candidate questions were generated with LLM assistance using study-specific information and BixBench-inspired few-shot examples, followed by manual prompt refinement. Questions underwent manual review for consistency with the intended analysis process, deduplication and disambiguation, ground-truth calibration, and balance of task complexity and type, yielding 26 questions across ten GWAS datasets. **b**, Agent execution and evaluation strategy. Agents received the benchmark question together with the corresponding GWAS summary statistics and generated free-text responses and execution traces. Each question was executed independently five times. Each run was evaluated separately under each of the three LLM-as-a-judge protocols. In open-answer evaluation, the judge compared the question, ground truth and agent free-text response and assigned a binary score. In multiple-choice evaluation with refusal, the judge mapped the initial agent response and execution trace to options A–D or E (insufficient information). In multiple-choice evaluation without refusal, the judge was required to map the response and execution trace to one of options A–D.

**Extended Data Fig. 3.**
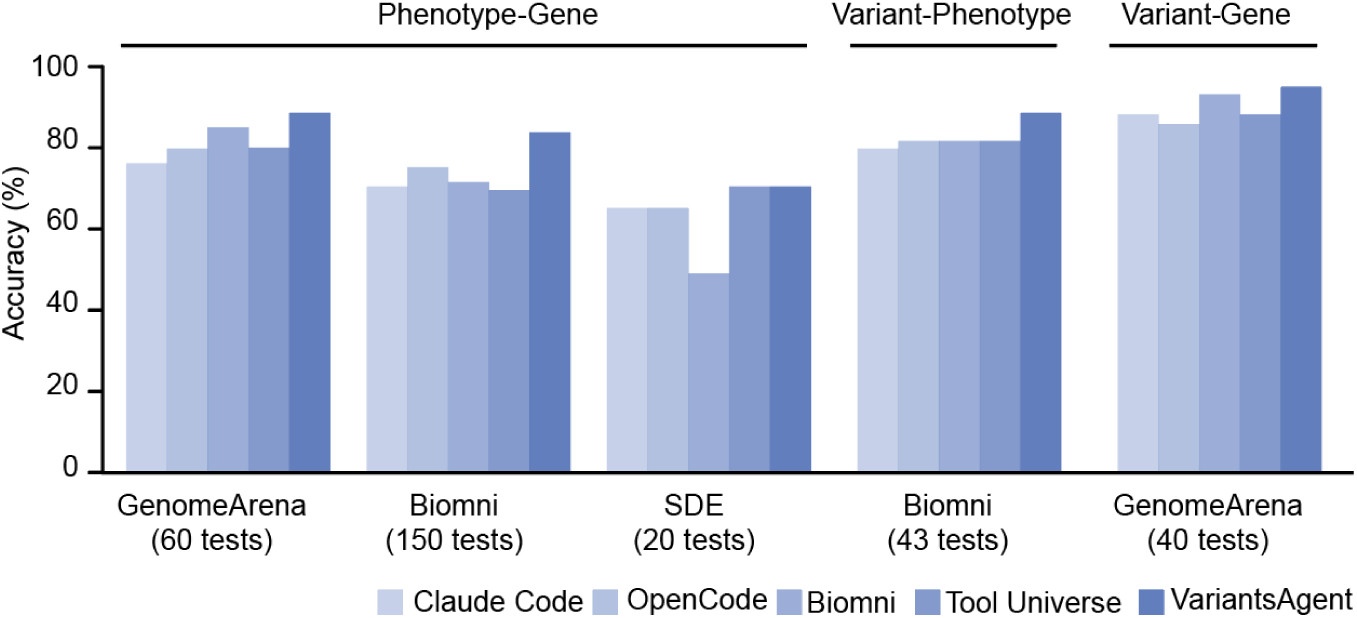
Performance across published genetic-reasoning benchmark classes. Accuracy of Claude Code, OpenCode, Biomni, Tool Universe and VariantAgent across published phenotype–gene, variant–phenotype and variant–gene benchmarks. Phenotype–gene evaluation comprised GenomeArena (60 tests), Biomni (150 tests) and SDE (20 tests); variant–phenotype evaluation comprised 43 Biomni tests; and variant–gene evaluation comprised 40 GenomeArena tests. Bars show accuracy for each available agent or baseline within each dataset.

**Extended Data Fig. 4.**
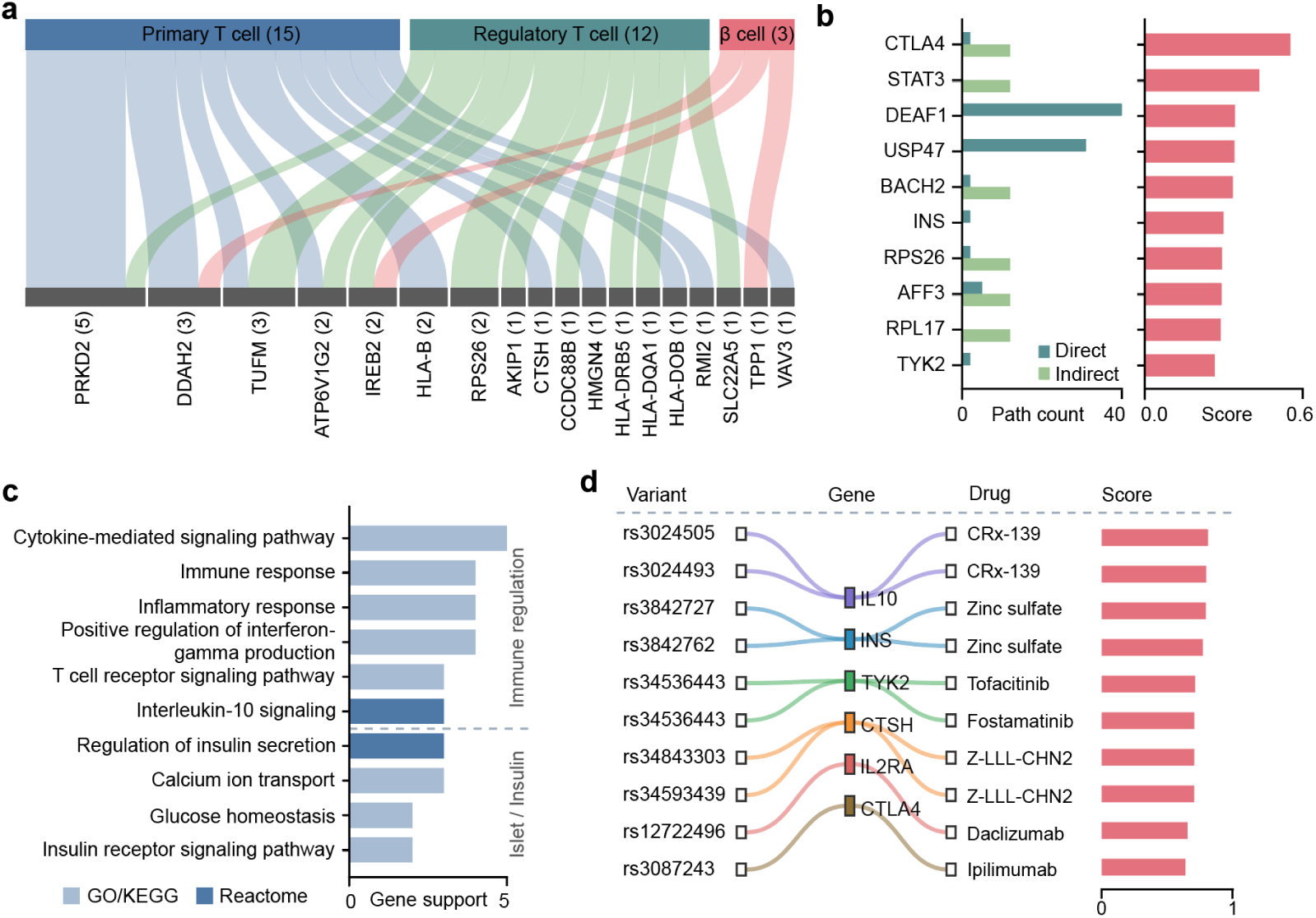
Experimental and knowledge-graph support for T1D gene, pathway and translational prioritization. **a**, Cell-type-specific CRISPR perturbation evidence for prioritized T1D candidate genes. CRISPR evidence was aggregated across high T1D-context cell types after deduplication of individual screens. Numbers in parentheses above each cell-type group indicate the cumulative edge-weight support across high-context CRISPR evidence records for that cell type (for example, Primary T cell, 15). Numbers in parentheses below individual genes indicate the number of deduplicated CRISPR screens supporting that gene across the displayed high-context cell types (for example, *PRKD2*, 5). Flow width reflects the corresponding accumulated support. **b**, Knowledge-graph-supported prioritization of T1D candidate genes. Direct support denotes edges directly connecting the T1D trait node to a gene, whereas indirect support denotes multi-hop paths of two or more edges connecting T1D to the gene. Left, numbers of direct and indirect supporting paths for each candidate gene. Right, the corresponding composite prioritization score, integrating graph-based and evidence attributes as defined in Methods. **c**, Functional programs supported by prioritized genes, ranked by the number of supporting genes. Terms are grouped into immune-regulatory and islet/insulin-related processes. Colours indicate pathway annotation source, with green representing GO/KEGG and blue representing Reactome. **d**, Ranked variant–gene–drug/chemical relationships supported by integrated post-GWAS and knowledge-graph evidence. Sankey links connect representative variants to candidate genes and downstream drug or chemical entities; bars show the corresponding path scores. These relationships represent translational evidence paths rather than clinical treatment recommendations.

**Extended Data Fig. 5.**
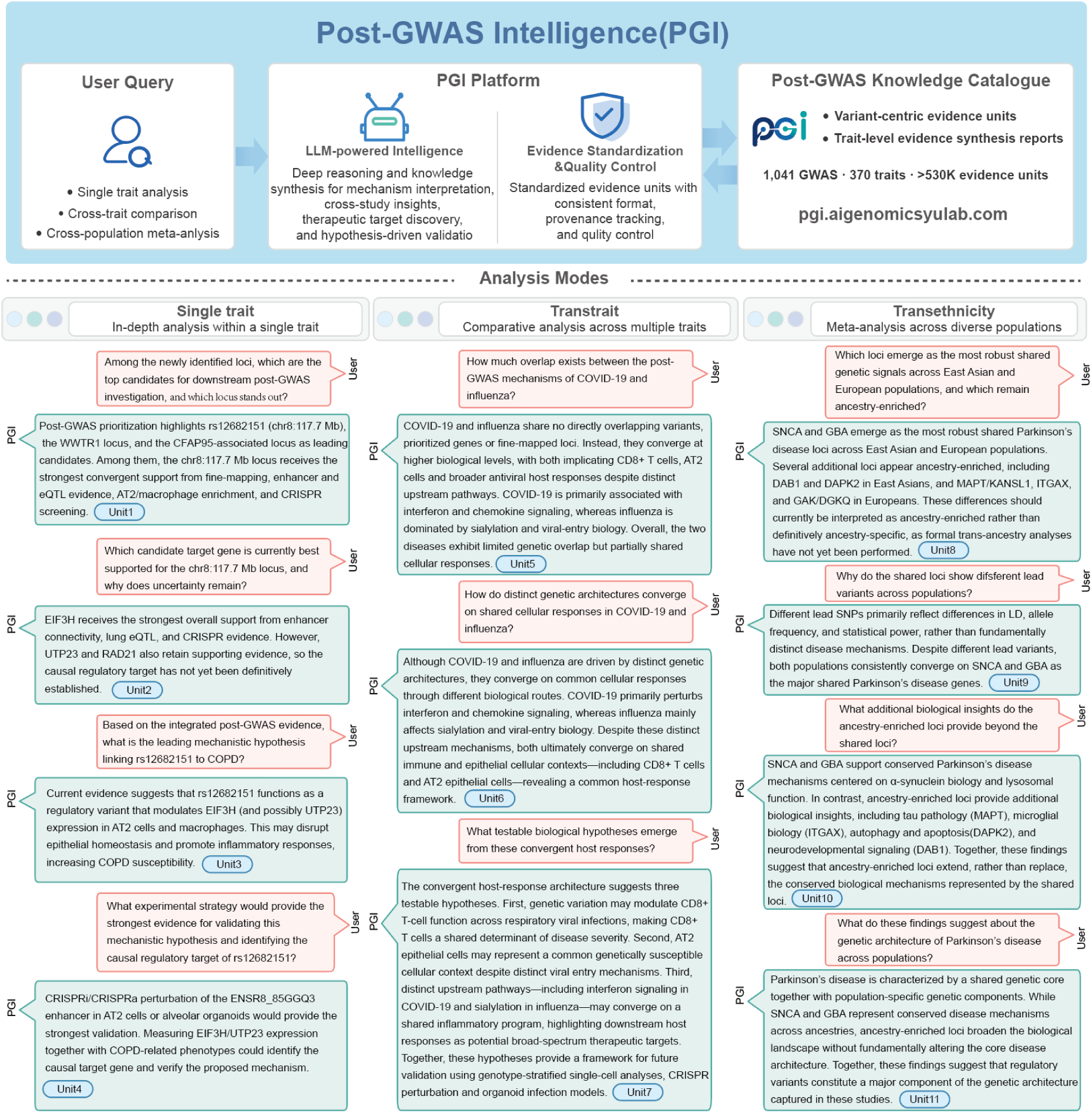
Evidence-grounded conversational interrogation of the PGI post-GWAS knowledge catalogue. The PGI conversational interface provides natural-language access to variant-centred evidence units and trait-level evidence synthesis reports generated from 1,041 GWAS across 370 harmonized traits. Queries are grounded in retrieved catalogue evidence and preserve links to the underlying analytical provenance. Three representative interaction modes are shown. Left, single-trait analysis: iterative interrogation of a COPD locus progresses from candidate prioritization to target-gene assessment, mechanistic hypothesis generation and proposal of an experimental validation strategy. Middle, cross-trait analysis: comparison of COVID-19 and influenza illustrates how distinct genetic architectures can be examined for convergence at cellular and mechanistic levels and translated into testable hypotheses. Right, trans-ethnic analysis: comparison of Parkinson’s disease across East Asian and European populations distinguishes shared loci from ancestry-enriched signals and interprets differences in the context of population-specific genetic architecture. The interface supports evidence retrieval and synthesis but does not add new evidence to the underlying catalogue.

